# Modification of the antibiotic, colistin, with dextrin causes enhanced cytotoxicity and triggers apoptosis in myeloid leukemia

**DOI:** 10.1101/2023.11.09.565276

**Authors:** Siân Rizzo, Mathieu Varache, Edward J. Sayers, Arwyn T. Jones, Alex Tonks, David W. Thomas, Elaine L. Ferguson

## Abstract

Acute myeloid leukemia (AML) remains difficult to treat due to its heterogeneity in molecular landscape, epigenetics and cell signaling alterations. Precision medicine is a major goal in AML therapy towards developing agents that can be used to treat patients with different ‘subtypes’ in combination with current chemotherapies. We have previously developed dextrin-colistin conjugates to combat the rise in multi-drug resistant bacterial infections and overcome dose-limiting nephrotoxicity. Recent evidence of colistin’s anticancer activity, mediated through inhibition of intracellular lysine-specific histone demethylase 1 (LSD1/KDM1A), suggests that dextrin-colistin conjugates could be used to treat cancer cells, including AML. This study aimed to evaluate whether dextrin conjugation (which reduces *in vivo* toxicity and prolongs plasma half-life) could enhance colistin’s cytotoxic effects in myeloid leukemia cell lines and compare the intracellular uptake and localization of the free and conjugated antibiotic. Our results identified a conjugate (containing 8,000 g/mol dextrin with 1 mol% succinoylation) that caused significantly increased toxicity in myeloid leukemia cells, compared to free colistin. Dextrin conjugation altered the mechanism of cell death by colistin, from necrosis to caspase 3/7-dependent apoptosis. In contrast, conjugation via a reversible ester linker, instead of an amide, had no effect on the mechanism of the colistin-induced cell death. Live cell confocal microscopy of fluorescently-labelled compounds showed both free and dextrin-conjugated colistin were endocytosed and co-localized in lysosomes and increasing the degree of modification by succinoylation of dextrin significantly reduced colistin internalization. Whilst clinical translation of dextrin-colistin conjugates for the treatment of AML is unlikely due to the potential to promote AMR and the relatively high colistin concentrations required for anticancer activity, the ability to potentiate the effectiveness of an anticancer drug by polymer conjugation, while reducing side effects and improving biodistribution of the drug, is very attractive, and this approach warrants further investigation.

**Graphical abstract:** 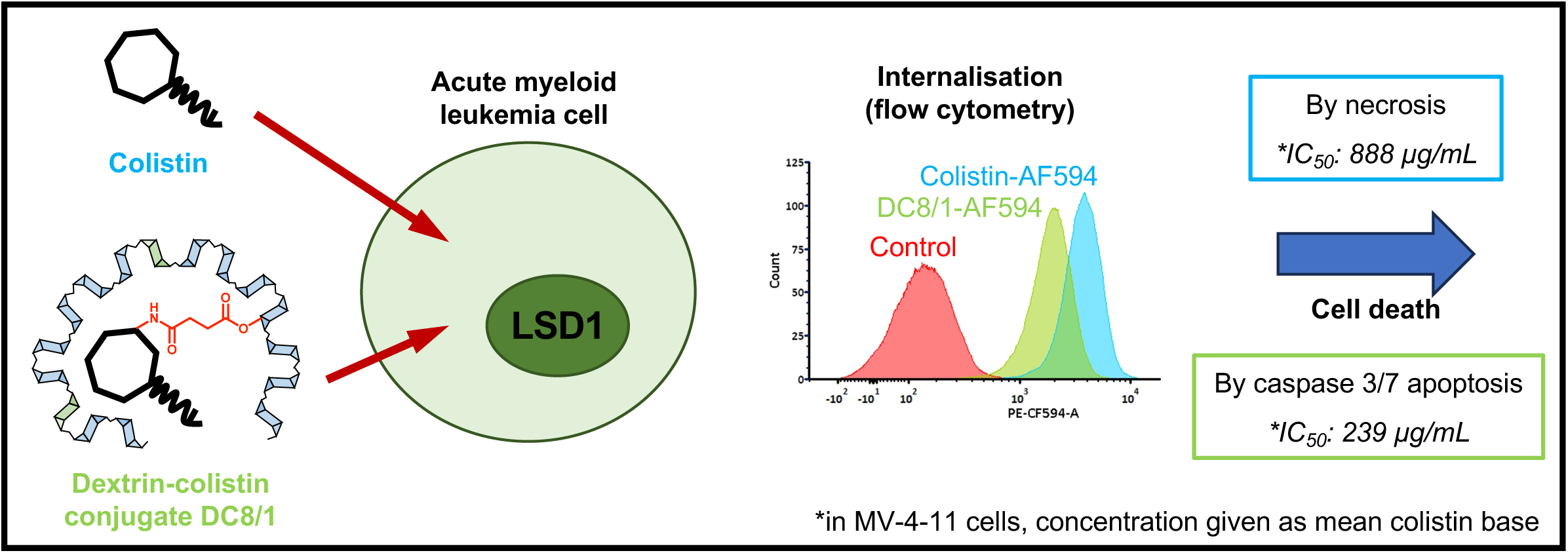

## Introduction

Acute myeloid leukemia (AML) is characterized by a block in hemopoietic differentiation where the hemopoeitic stem cell (HSC) fails to develop into mature myeloid cells resulting in bone marrow failure.^1^ AML arises due to a wide range of molecular abnormalities and genetic mutations, making the disease difficult to treat.^2^ Although AML accounts for fewer than 1% of all new cancer cases,^3^ recurrence after complete remission is common. Importantly, infection, due to treatment-related immunosuppression, expression of immunosuppressive molecules or specific defects in the immune response, remains a major cause of death in patients with AML, requiring antibacterial or antiviral therapies to treat infections during first-line treatment.^4^ Given the high costs, high risk of failure and protracted development times of new chemotherapy drugs, exploiting the off-target effects of existing antimicrobial agents is an attractive approach to improve survival rates in cancer p[atients.

Polymer therapeutics are an important class of drugs that use conjugation of a water-soluble polymers to improve the therapeutic index of drugs, including for oncology. Oncaspar^®^, a PEG-L-asparaginase conjugate was one of the first polymer therapeutics to receive FDA approval, for the treatment of acute lymphoblastic leukemia (ALL).^5^ There are currently several nanomedicines in clinical use or undergoing clinical trials for the treatment of AML, including liposomes (e.g. VYXEOS^®^; liposomal daunorubicin and cytarabine),^6^ polymeric nanoparticles (e.g. AZD2811; aurora kinase B inhibitor-loaded Accurin^TM^ polymeric nanoparticles)^7^ and polymer-drug conjugates (e.g. N- (2-hydroxypropyl) methacrylamide (HPMA) copolymer-cytarabine and GDC-0980 conjugates).^8^ We have previously developed bio-triggered polymer therapeutics, based on the attachment of the polysaccharide, dextrin, to a bioactive molecule, including phospholipase A2 (PLA2),^9^ epidermal growth factor^10^ and colistin (also known as polymyxin E),^11^ as treatments for breast cancer, chronic wounds and bacterial infection, respectively.

Recently, it has been shown that the polymyxin antibiotics, polymyxin B and colistin, can kill some types of cancer cells,^12^ including the ALL cell line REH.^13^ Polymyxin’s anticancer activity is believed to be mediated through inhibition of lysine-specific histone demethylase 1 (LSD1, also referred to as lysine-specific demethylase 1A: KMD1A),^14^ a nuclear histone-modifying enzyme which is overexpressed in AML leading to poorer clinical outcomes.^15^ Recently, it has been shown that inhibition of LSD1/KDM1A activity results in inhibition of AML cell proliferation and enhanced differentiation.^15^ Thus, several LSD1/KDM1A inhibitors, including GSK2879552 and tranylcypromine, are currently in clinical trials for cancer treatment,^16^ including AML, as a combination therapy.^17^ Given the efficacy of colistin as a prophylactic treatment of neutropenia in AML,^18^ polymyxin antibiotics have the potential to simultaneously act as antibacterial and anticancer agents. However, dose-limiting neuro-and nephrotoxicity prevent their routine use,^19^ which may be overcome by polymer conjugation, as we have shown in our previous work, where dextrin-colistin conjugates reduced toxicity *in vitro* and *in vivo*, and extended plasma half-life.^11,20,21^ Our recent studies have demonstrated that conjugation of dextrin to colistin reduced cellular internalization of the antibiotic by kidney cells and caused reduced cytotoxicity.^22^ In parallel, dextrin conjugation to colistin dramatically diminished both proximal tubular injury and renal accumulation of colistin in mice receiving twice-daily doses of the antibiotic. Therefore, we hypothesized that dextrin-colistin conjugates may be repurposed as an anticancer therapy, offering improved biodistribution, sustained drug exposure and reduced side-effects.

Polymer conjugation has been widely used as a means of improving drug delivery.^23,24^ However, while polymer-protein and -peptide conjugation has shown particular success for their delivery to extracellular targets, intracellular delivery has proved to be more challenging.^25^ Our previous studies have shown that dextrin conjugation has a varying effect on intracellular uptake of a drug; while attachment of dextrin to PLA2 significantly increased the proportion of internalized PLA2 by MCF-7 breast cancer cells,^26^ less colistin internalization in HK-2 proximal tubule cells was observed when conjugated to dextrin.^22^ Therefore, it was important to establish whether conjugation of colistin to dextrin would alter its endocytic properties in AML cells.

The aim of this study was to investigate and quantify the intracellular delivery of 4 different dextrin-colistin conjugates (containing 8,000 or 51,000 g/mol dextrins, having 1 or 10 mol% succinoylation) to demonstrate whether dextrin conjugation can enhance colistin’s anticancer activity. To achieve this, *in vitro* cytotoxicity, cellular uptake, and localization of fluorescently labelled conjugates were assessed in three leukemia cell lines.

## Material and methods

### Materials

Type I dextrin prepared from corn (Mw = 8,000 g/mol, degree of polymerization (DP = 50), colistin sulfate, N-hydroxysulfosuccinimide (sulfo-NHS), dimethyl sulfoxide (DMSO), dimethylformamide (DMF), Hoechst 33342 solution and staurosporine were purchased from Sigma-Aldrich (Poole, UK). Dextrin (Mw 51,000 g/mol, DP = 315) was from ML Laboratories (Liverpool, UK). Disodium hydrogen phosphate, potassium dihydrogen phosphate, potassium chloride, 4- dimethylaminopyridine (DMAP), BCA protein assay kit, N,N_′_-dicyclohexyl carbodiimide (DCC), 1- ethyl-3-(3-(dimethylamino)propyl carbodiimide hydrochloride) (EDC), AlexaFluor^®^ 594 (AF594) cadaverine, AF594 succinimidyl ester, sodium chloride, Gibco^TM^-branded keratinocyte serum-free medium (K-SFM) with L-glutamine, epidermal growth factor (EGF), bovine pituitary extract (BPE), 0.05% w/v trypsin-0.53 mM EDTA, Iscove’s modified Dulbecco’s medium (IMDM) with GlutaMAX^TM^, fetal bovine serum (FBS) and Roswell Park Memorial Institute (RPMI) 1640 medium with GlutaMAX^TM^ were obtained from ThermoFisher Scientific (Loughborough, UK). CellTiter-Blue® (CTB) cell viability assay kit, CytoTox-One^TM^ homogeneous membrane integrity assay kit and Caspase-Glo® 3/7 assay system kit were from Promega, WI, USA. Recombinant human granulocyte-monocyte colony stimulating factor (rhGM-CSF) was from Peprotech (London, UK). Pullulan gel filtration standards (M_w_ = 180−788,000 g/mol) were purchased from Polymer Laboratories (Church Stretton, UK) and Shodex (Tokyo, JP). Unless otherwise stated, all chemicals were of analytical grade and used as received. All solvents were of general reagent grade (unless stated) and were from Fisher Scientific (Loughborough, UK).

### Synthesis and characterization of dextrin-colistin conjugates and AF594-labelled probes

Amide-linked dextrin–colistin conjugates, containing dextrin (*M*w = 8,000 or 51,000 g/mol) with 1 and 10 mol% succinoylation, were synthesized using EDC and sulfo-NHS and characterized as described previously.^11^ To synthesize ester (E)-linked conjugates, succinoylated dextrins (1, 2.5 and 10 mol% succinoylation) were conjugated to colistin using DCC and DMAP. Briefly, for 1 mol% succinoylated dextrin, succinoylated dextrin (1000 mg, 0.125 mmol), DCC (25.5 mg, 0.25 mmol) and DMAP (7.5 mg, 0.125 mmol) were dissolved under stirring in anhydrous DMSO (10 mL) in a 50 mL round-bottomed flask, and left stirring at 21°C for 15 min. Subsequently, colistin sulphate (176 mg, 0.25 mmol) was added, and the mixture was left stirring overnight at 21°C. To stop the reaction, the mixture was poured into excess chloroform (∼100 mL). Resulting precipitates were isolated by filtration and dissolved in ultrapure water (H_2_O; Milli-Q^®^ filtered to 18.2 M_Ω_ cm^-1^) (10 mL), then stored at −20°C before purification by fast protein liquid chromatography (FPLC) (AKTA Purifier; GE Healthcare, UK) using a pre-packed HiLoad Superdex 30 26/600 column equipped with a UV detector, using Unicorn 5.31 software (GE Healthcare, Amersham, UK) for data analysis.

To enable visualization of conjugates by flow cytometry and confocal microscopy, succinoylated dextrin, colistin and dextrin-colistin conjugate were fluorescently labelled with AlexaFluor^®^ 594 (AF594) and characterized according to previously published methods,^27–29^ including spectrophotometric and fluorometric analysis. Briefly, to prepare AF594-labelled colistin, the antibiotic (16.5 mg) was solubilized under stirring in PBS (1 mL, pH 7.4) in a 10 mL round-bottomed flask. AF594 succinimidyl ester (9.5 mg, from a stock solution of 10 mg/mL in anhydrous DMF, stored at -20°C until use) was added dropwise. Then, the reaction mixture was stirred at room temperature, protected from light. After 1 h, the solution was purified by FPLC (as above) using a prepacked Superdex 30 26/600 column coupled with a UV detector set at 210, 280 and 550 nm. The reaction mixture (5 mL) was injected into a 5 mL loop and eluted using 0.1 M ammonium acetate (pH 6.9, 0.22 µm filter-sterilized) at a flow rate of 2.5 mL/min. Fractions (15 mL) containing colistin-AF594 (typically between 230 and 290 mL) were identified by HPLC-fluorescence, then pooled and desalted by freeze-drying (×5) to remove ammonium acetate. The final compound was stored at -20°C until use. To prepare AF594-labelled dextrin, succinoylated dextrin (160 mg, 8,000 g/mol, 10 mol%) was dissolved under stirring in PBS buffer (5 mL, pH 7.4) in a 10 mL round-bottomed flask. To this, EDC (18.9 mg) and sulfo-NHS (21.4 mg) were added, and the mixture stirred for 15 min before addition of AF594 cadaverine (8 mg, from a stock solution of 10 mg/mL in anhydrous DMF, stored at -20°C until use). The reaction mixture was stirred in the dark for 5 h prior to purification by size exclusion chromatography (SEC, disposable PD-10 desalting column containing Sephadex G-25). To prepare the AF594-labelled dextrin-colistin conjugate with 10 mol% succinoylation, a dextrin-AF594 intermediate was prepared, as described above, conjugated to colistin and purified by FPLC, as previously described.^11^ As 1 mol% succinoylated 8,000 g/mol dextrin would not contain sufficient reactive groups to attach both, colistin and AF594, to prepare AF594-labelled dextrin-colistin conjugate with 1 mol% succinoylation, a colistin-AF594 intermediate was initially prepared, as above, and subsequently conjugated to 1 mol% succinoylated dextrin and purified as previously described^11^.

The total AF594 content of AF594-labelled dextrin and dextrin-colistin conjugates was determined by measuring absorbance at 485 nm. Free AF594 content was assessed by measuring fluorescence (_λex_ = 588 nm, _λem_ = 612 nm, gain 1000) of fractions (1 mL) eluting from a PD-10 column, according to previously described methods.^29^ Analysis of colistin-AF594 was performed using LC-MS on a Synapt G2-Si quadrupole time-of-flight (QTOF) mass spectrometer (Waters, U.K.), operating in the positive electrospray ionization mode, coupled to an ACQUITY H-Class UPLC system (Waters, Wilmslow, UK). Separation was accomplished using an ACQUITY UPLC BEH column (1.7 μm, 2.1 x 100 mm, Waters) inside a column oven at 40°C. A multistep gradient method was used (0-2 min, 98% A; 2-20 min, 2% A; flow rate 0.3 mL/min), where mobile phase A is water (0.1% formic acid), and mobile phase B is acetonitrile (0.1% formic acid). Analysis of the purity of AF594-labelled dextrin and dextrin-colistin conjugates was performed using PD-10 columns (Cytiva, Little Chalfont, UK), by analysis of SEC fractions for fluorescence and protein content (by BCA reagent).

### Hemolytic activity

All animal experiments were conducted according to the United Kingdom Use of Animals (Scientific Procedures) Act 1986. Animal work was reviewed by the Animal Welfare and Ethical Review Body under the Establishment License held by Cardiff University and authorized by the UK Home Office. Fresh blood was extracted from recently euthanized male Wistar rats (∼250 g body weight) by cardiac puncture and added to 4 mL PBS (pH 7.4) in a heparin/lithium blood tube. The tube was centrifuged at 400 x *g* for 10 min at 4°C to extract red blood cells (RBCs), then washed a further two times. Following the final wash, the RBC pellet was diluted to 2% w/v with PBS. Subsequently, this diluted RBC suspension was added to a 96-well plate (100 μL/well, replicates n=6) containing an equal volume of test compounds, PBS (negative control) or Triton X-100 (1% v/v) (positive control). Following incubation for 1 h at 37°C, the plate was centrifuged at 400 x *g* for 10 min at 4°C. The supernatant (100 μL) of each well was transferred to a 96-well plate, and absorbance at 550 nm was read using a Fluostar Omega microtiter plate reader. Cells were plated with 6 technical replicates per plate (n=6) and each experiment was performed three times. The negative control (PBS) absorbance was subtracted, and the results were expressed as mean percentage of maximum (Triton X-100) hemoglobin released ± 1SD (n=6).

### Cell culture

Human kidney proximal tubule cells (HK-2) and the leukemia cell lines MV-4-11 and TF-1 were obtained from ATCC (Manassas, USA). The leukemia cell line THP-1 was from ECACC (UK Health Security Agency). Characteristics of the cell lines used are shown in Table S1. Cells were screened to be free of mycoplasma contamination upon thawing and monthly thereafter using a Venor GeM Classic Mycoplasma Detection Kit from Minerva Biolabs (Berlin, Germany). Cell lines were maintained in log-phase proliferation at 37°C with 5% CO_2_/air and cultured in their respective culture medium (Table S1). HK-2 cells were passaged using 0.05% w/v trypsin-0.53 mM EDTA.^30^

### Evaluation of *in vitro* cytotoxicity

Cytotoxicity of colistin sulfate and the dextrin-colistin conjugates was assessed in the above cell lines using a multiplexed assay system, to measure cell viability, membrane integrity (lactate dehydrogenase (LDH) release), caspase activity/apoptosis and DNA content as described previously.^31^ In summary, cells were seeded into sterile black, clear-based 96-well microplates (HK- 2 at 2,500 cells/ well: all other cell lines at 10,000 cells/ well in 0.1 mL of complete media). Cultures were incubated at 37°C with 5% CO_2_ for 1 h (MV-4-11, THP-1, TF-1) or, to allow adherent cells to adhere, for 24 h (HK-2). Test compound stock solutions were prepared in PBS (0.22 μm filter-sterilized) and used to supplement the complete media. Test compounds were evaluated in triplicate at concentrations up to 1 mg/mL colistin base with respective vehicle-only control (PBS), while 1-10 µM staurosporine (apoptosis) and 100 µM Triton-X100 (necrosis) were used as positive controls.

Following a further 24 or 72 h incubation, plates were processed as follows, protected from light throughout. To measure necrosis, 25 μL of supernatant from each well was transferred to a fresh black 96-well microplate, containing 25 μL of CytoTox-One^TM^ assay reagent. Cultures were gently mixed then incubated at 21°C for 10 min, then a “stop solution” was added and fluorescence measured at _λex_ = 560 nm, _λem_ = 590 nm using a Fluostar Omega microplate reader. Total cellular LDH activity was measured after the addition of LDH lysis solution to cells, prior to addition of the assay reagent. Next, to measure cytotoxicity and DNA content, CTB reagent was supplemented with Hoechst 33342 (100 µg/mL) and added to the wells of the original microplate (10 µL/well). Cultures were gently mixed followed by incubation at 37°C with 5% CO_2_ in air for 1 h and fluorescence measured at _λex_ = 560 nm, _λem_ = 590 nm and _λex_ = 340 nm, _λem_ = 460 nm (for cytotoxicity and DNA content, respectively) using a Fluostar Omega microplate reader. Finally, 60 μL of Caspase-Glo 3/7 assay reagent was added to each well of the original microplate. Plates were gently agitated then incubated at 20°C with 5% CO_2_ in air for 1 h, followed by luminescence assays using a Fluostar Omega microplate reader. Cells were plated with 3 technical replicates per plate (n=3) and each experiment was performed twice. Data was corrected for no-cell background, then expressed as mean percentage of the response of vehicle-only control cells ± 1SD (n=3). Relative IC_50_ values were determined using the non-linear regression analysis of dose-response-inhibition using a 4-parameter logic model in GraphPad Prism (version 9.3.1, 2021; San Diego, CA, USA).

To monitor apoptosis and necrosis continuously over 72 h, a RealTime-Glo™ Annexin V apoptosis and necrosis assay (Promega, Southampton, UK) was used. Briefly, MV-4-11 cells were plated into sterile black, clear-based 96-well microplates (4,500 cells/ well in 45 µL of complete media). Test compound solutions (10 µL) and 2x detection reagent (55 µL) were added to each well, in triplicate, and the plate was gently agitated before placing inside a Fluostar Omega plate reader (maintained at 37°C with 5% CO_2_). Fluorescence (_λex_ = 485 nm, _λem_ = 525-530 nm) and luminescence were measured at regular timepoints for up to 72 h. Data was corrected for no-cell background and expressed as a percentage of the response of vehicle only control cells. Staurosporine and Triton-X100 were used as positive controls for apoptosis and necrosis, respectively, as described above. Cells were plated with 3 technical replicates per plate (n=3) and each experiment was performed once.

### Cell cycle analysis

Cell cycle analysis of MV-4-11 cells following incubation with colistin sulfate, DC8/1 or DC8/10 at the previously determined IC_50_ value was performed according to a modified version of Ormerod’s protocol.^32^ Briefly, 500 µL cells at 10^5^ cell/ mL per well of a 24-well plate were incubated in complete media supplemented with test compounds (in duplicate) at their IC_50_ value at 37°C with 5% CO_2_ for 24 h. The contents of each well were diluted with 3 mL PBS, then centrifuged (350 x g for 5 min at 20°C), the cell pellet was resuspended in ice-cold PBS then centrifuged again prior to removing the supernatant. Subsequently, ice-cold 70% v/v ethanol was added dropwise under vortex to resuspend the cells, then incubated on ice. Following 30 min, cells were centrifuged (450 x g for 10 min at 4°C) then washed twice with ice-cold PBS, vigorously resuspending the pellet each time. To the cell pellet, 50 µL of RNase (100 µg/mL) solution was added then incubated at 37°C for 15 min. Finally, 200 µL of propidium iodide (PI; 50 µg/mL) solution was added to each tube 20 min prior to data acquisition with a Becton Dickinson FACS CANTO II Cell Analyser flow cytometer equipped with a 488 nm blue laser and 584/42 nm emission filter. Data were collected for 25,000 events per sample and data analyzed using FlowJo™ software, v10.8.1. MV-4-11 cells incubated with medium only were used to determine background fluorescence. Cell fragments, clumps and debris were excluded using sequential gating on a forward-scatter (FSC)-height vs side-scatter (SSC)-height cytogram and a DNA (FL2)-area vs DNA (FL2)-width cytogram, then the remaining single cells were displayed in a histogram of DNA-area (FL2), as described by Ormerod; the Watson (pragmatic) algorithm, a univariate cell cycle model, was employed to assess the proportion of cells in each cell cycle phase: G1, S and G2(M)^33^ using FlowJo™ software (version 10.8.1). Cells were plated with 2 technical replicates per plate (n=2) and each experiment was performed four times.

### Colony formation

Colony-forming ability of MV-4-11 and THP-1 cells following incubation with colistin sulfate, DC8/1 or DC8/10 at the previously determined IC_50_ value was measured using a human Colony Forming Cell (CFC) assay in methylcellulose-based media, according to manufacturer’s instructions (R&D Systems, Abingdon, UK). Briefly, cells (10^5^ cells/mL) were incubated in complete media supplemented with 10% v/v test compounds at their IC_50_ value or with vehicle (PBS) at 37°C with 5% CO_2_ for 24 h. Then, the equivalent of 667 THP-1 or 400 MV-4-11 cells were added in duplicate to 1 mL human methylcellulose base media supplemented with recombinant human granulocyte-monocyte colony stimulating factor (10 ng/mL), interleukin-3 (10 ng/mL) and stem cell factor (20 ng/mL) in a 35 mm culture dish (Nunclon, Fisher Scientific). Following incubation at 37°C with 5% CO_2_ for 10 days, colonies containing >32 cells were enumerated by inverted light microscopy. The results were expressed as mean percentage colony-forming units (CFU) of vehicle-only control (n=2).

### Determination of cellular uptake

To investigate cell association at 4 or 37°C, leukemia cells were first resuspended at 200,000 cells/well of a 96-well plate in 75 µL complete medium (without phenol red). Experiments at 37°C were conducted with standard cell culture conditions, but for low temperature experiments, cells were pre-incubated for 30 min at 4°C prior to the addition of the probe. Solutions of fluorescent probes were freshly prepared in complete medium at sub-toxic, equivalent concentrations of AF594 base (1μg AF594 base/mL), filter-sterilized (0.22 μm), then equilibrated to either 37 or 4°C for 30 min. Probe solutions were added to each well (75 µL) containing cells and incubated at 4 or 37°C for 2 h. Subsequently, plates were placed on ice before transferring the cell suspension into individual tubes and washing twice by centrifugation (350 x *g*, 5 min) with ice-cold PBS (2 x 3 mL). Finally, cells were resuspended in ice-cold PBS (200 μL) prior to data acquisition using a Becton Dickinson LSR Fortessa Cell Analyser flow cytometer equipped with a yellow-green laser (561 nm) and emission filter for 585/15 nm. For each sample.10,000 events were collected and analyzed using FlowJo™ software v 10.8.1. Control cells incubated with medium only were used to determine background fluorescence. Throughout, results were corrected for cell autofluorescence and expressed as (geometric mean × % positive cells)/100, where % positive cells was calculated as 100 – % cells in M1 (see Figure 5a-d for example of gating). Internalization was calculated by subtracting the cell-associated fluorescence at 4°C (extracellular binding) from that at 37°C (intracellular uptake plus extracellular binding) and expressed as mean ± SD. All uptake studies were performed using colistin concentrations that have previously been shown to be non-toxic. Cells were plated with three technical replicates per plate (n=3); each experiment was performed three times. Results are expressed as mean fluorescence ± 1SD (n=3).

### Intracellular localization

Cells were incubated in complete media (without phenol red, at 6.67 x 10^5^ cells/mL) containing dextran-AF488 (150 µg/mL) at 37°C with 5% CO_2_ /in air. After 3 h, excess dextran-AF488 was removed by addition of complete media (10x labelling volume) followed by centrifugation at 350 x *g* for 5 min. The cell pellet was then resuspended in complete media (10^5^ cells/150 µL) with and without AF594 probes (5 µg AF594 base/mL) and incubated at 37°C with 5% CO_2_ for 2 h in 96-well plates. In all cases, colistin concentration was below IC_50_. The cells were washed twice with 10x volume complete medium, as described above. After the final wash, 10^5^ cells/mL cells were cultured in fresh medium at 37°C with 5% CO_2_ for a further 2 h (short chase) or overnight (16 h, long chase). Control cells (not incubated with a probe) were used to account for autofluorescence.

Confocal laser scanning microscopy was performed using a Leica SP5 system (37°C, 5% CO_2_). Confocal imaging was performed sequentially with the 405 nm (Hoechst 33342) and 543 nm (AF594) lasers captured concurrently and 488 nm (AF488) laser captured sequentially between lines. Images were captured using a HCX PL APO CS 100.0×1.40 oil immersion objective at 400 Hz with a line average of 3 (bi-directional scanning), with the pinhole set to 1 airy unit. Images were acquired with a raster size of 1024 x 1024 and a zoom of 1.12 to give an apparent pixel size of 138 x 138 nm (XY). At least eight representative images (single section) were obtained from each sample; typical results are shown. Images were analyzed and processed using ImageJ.^34^

### Statistical analysis

GraphPad Prism (version 9.3.1, 2021; San Diego, CA, USA) software was used to perform statistical analyses. Statistical significance indicated by * *p* < 0.05, ** *p* < 0.01, *** *p* < 0.001 and **** *p* < 0.0001. One-way analysis of variance (ANOVA) was used to evaluate multiple group comparisons (*n* _≥_ 3) followed by Dunnett’s post hoc test to account for multiple comparisons, unless otherwise stated.

## Results

Amide-and ester-linked dextrin-colistin conjugates (containing 8,000 or 51,000 g/mol dextrins, having 1 or 10 mol% succinoylation) were synthesized to compare their *in vitro* cytotoxicity and colony forming ability of treated cells. Fluorescent dextrin, colistin and dextrin-colistin conjugate were subsequently obtained by chemical attachment of Oregon Green (OG) to enable measurement of cellular uptake and localization in leukemia cell lines.

### Synthesis and characterization of dextrin-colistin conjugates

The characteristics of dextrin-colistin conjugates used in these studies are summarized in Table 1. Free colistin content, analyzed by FPLC, was determined to be <3%.

**Table 1.**
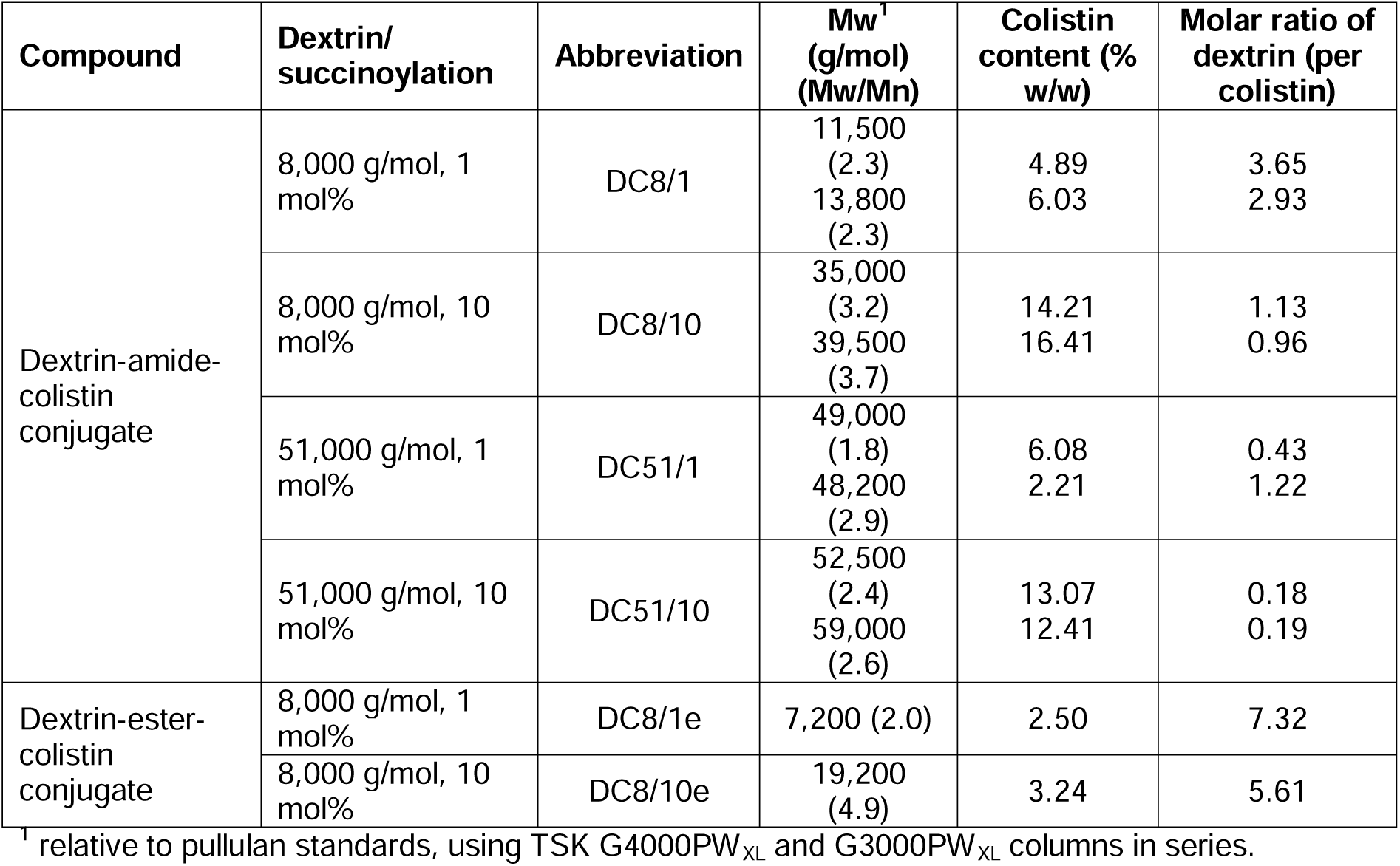
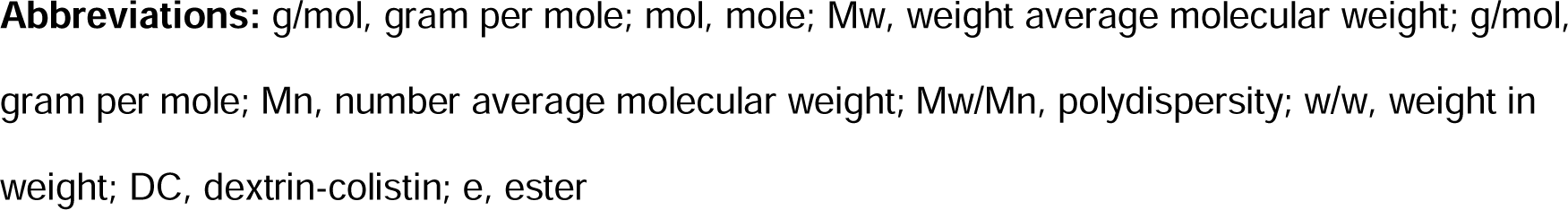
Summary of the properties of the dextrin-colistin conjugates used in this study.

| Compound | Dextrin/succinylation | Abbreviation | Mw<sup>†</sup><br>(g/mol)<br>(Mw/Mn) | Colistin<br>content (%<br>w/w) | Molar ratio of<br>dextrin (per<br>colistin) |
| --- | --- | --- | --- | --- | --- |
| Dextrin-amide-colistin conjugate | 8,000 g/mol, 1 mol% | DC8/1 | 11,500<br>(2.3)<br>13,800<br>(2.3) | 4.89<br>6.03 | 3.65<br>2.93 |
|  | 8,000 g/mol, 10 mol% | DC8/10 | 35,000<br>(3.2)<br>39,500<br>(3.7) | 14.21<br>16.41 | 1.13<br>0.96 |
|  | 51,000 g/mol, 1 mol% | DC51/1 | 49,000<br>(1.8)<br>48,200<br>(2.9) | 6.08<br>2.21 | 0.43<br>1.22 |
|  | 51,000 g/mol, 10 mol% | DC51/10 | 52,500<br>(2.4)<br>59,000<br>(2.6) | 13.07<br>12.41 | 0.18<br>0.19 |
| Dextrin-ester-colistin conjugate | 8,000 g/mol, 1 mol% | DC8/1e | 7,200 (2.0) | 2.50 | 7.32 |
|  | 8,000 g/mol, 10 mol% | DC8/10e | 19,200<br>(4.9) | 3.24 | 5.61 |

<sup>†</sup> relative to pullulan standards, using TSK G4000PW<sub>XL</sub> and G3000PW<sub>XL</sub> columns in series.
**Abbreviations:** g/mol, gram per mole; mol, mole; Mw, weight average molecular weight; g/mol, gram per mole; Mn, number average molecular weight; Mw/Mn, polydispersity; w/w, weight in weight; DC, dextrin-colistin; e, ester

### Hemolytic activity

The hemolytic activity of colistin and dextrin-colistin conjugates was measured with fresh rat RBCs to assess the effect of free and dextrin-conjugated colistin on normal blood cells. After exposure of RBCs to treatments for 24 h, only the highest dose of free colistin sulfate induced significant lysis, compared to vehicle alone (Figure 1). No hemolysis was observed for any of the dextrin-colistin conjugates up to 500 µg/mL. Control experiments with dextrin and succinoylated dextrins showed <10% hemolysis up to 10 mg/mL (data not shown).

**Figure 1.**
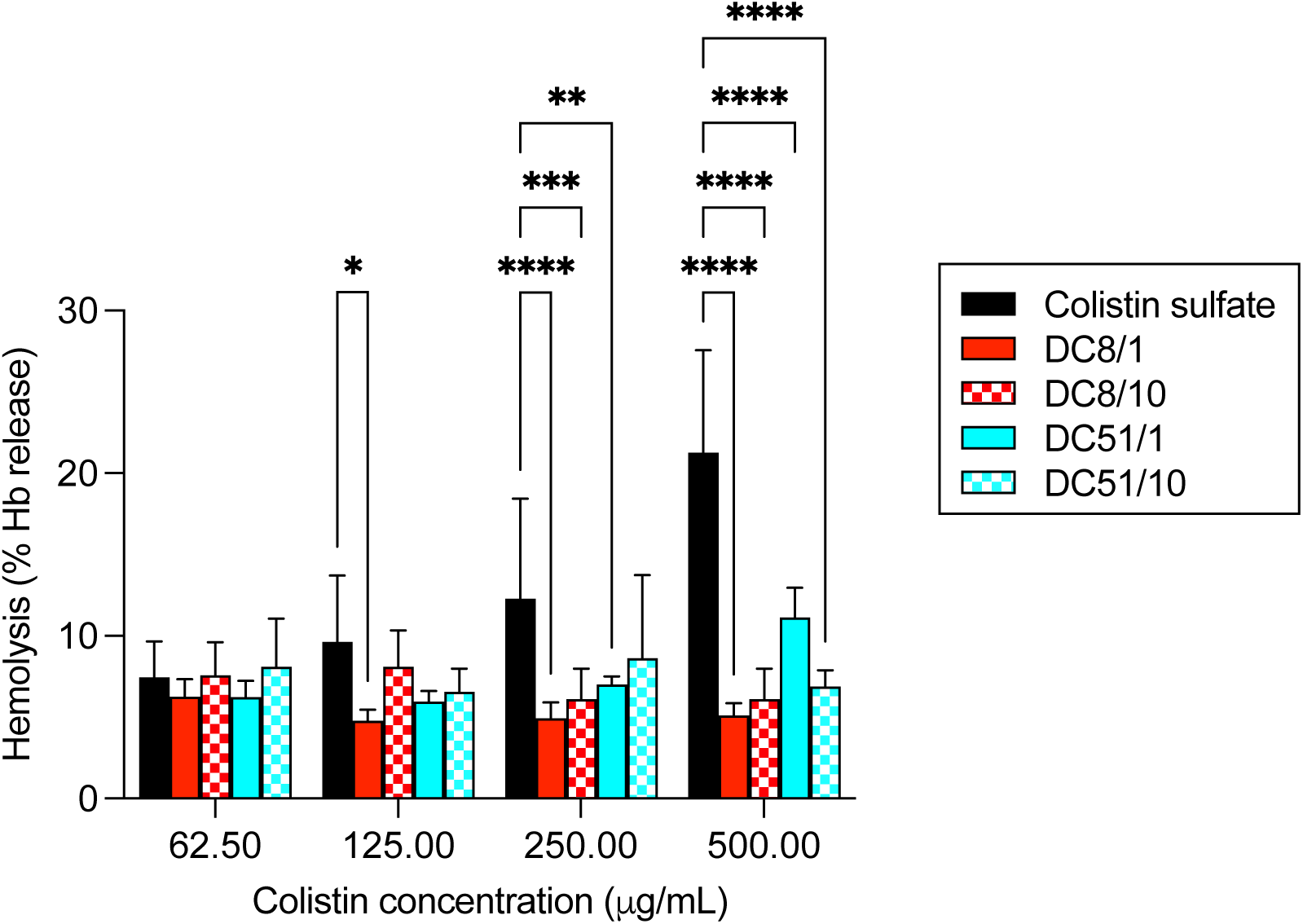
Hemolysis of rat erythrocytes following incubation for 24 h with colistin sulfate and dextrin-colistin conjugates at 37°C. Data is expressed as mean % Triton X-100 control ± 1SD, n = 6, where * indicates significance *p*<0.05, ** indicates significance *p*<0.01, *** indicates significance *p*<0.001 and **** indicates significance *p*<0.0001 compared to colistin sulfate. Where significance is not shown, *p*>0.05 (ns, not significant).

### *In vitro* cytotoxicity

Cytotoxicity was determined in three representative myeloid leukemia cells lines, in comparison to a non-cancerous human kidney proximal tubular cell line. As expected, cell viability decreased with longer incubation times in all cell lines. Where inhibition at 24 h achieved at least 50%, cell viability data was used to generate relative IC_50_ values, summarized in Table 2.

**Table 2.** Half maximal inhibitory concentration (IC_50_) values and fold-change (CTB assay) of colistin sulfate and dextrin-colistin conjugates after 24 h incubation.

| Compound | IC<sub>50</sub> (µg/mL colistin base, ± SD)<sup>1</sup> |  |  |  |
| --- | --- | --- | --- | --- |
|  | THP-1 | TF-1 | MV-4-11 | HK-2 |
| Colistin sulfate | 500 ± 89 | 670 ± 17 | 888 ± 37 | 447 ± 67 |
| DC8/1 | 396 ± 13 (0.8) | 315 ± 57 (0.5) | 239 ± 19 (0.3) | 653 ± 78 (1.5) |
| DC8/10 | >1000 | >1000 | >1000 | >1000 |
| DC51/1 | 724 ± 161 (1.5) | 819 ± 2 (1.2) | 880 ± 64 (1.0) | 638 ± 19 (1.4) |
| DC51/10 | >1000 | >1000 | >1000 | >1000 |

<sup>1</sup> Values in parentheses correspond to fold-change, compared to colistin sulfate.
**Abbreviations:** IC<sub>50</sub>, half-maximal inhibitory concentration; µg/mL, microgram per milligram; SD, standard deviation; DC, dextrin-colistin

Colistin sulfate exhibited greatest cytotoxicity against HK-2 cells (Figure 2a and S1a); amide-linked dextrin conjugation reduced toxicity of colistin in HK-2 by at least 1.4-fold, compared to the free drug, and was most effective in conjugates containing longer dextrin chains and higher degrees of succinoylation. Similarly, in the leukemia cell lines, conjugation to dextrin reduced cytotoxicity for three of the four amide-linked conjugates tested: DC8/10, DC51/1 and DC51/10 (Table 2, Figures 2e and S1e) by varying degrees. However, in all leukemia cell lines tested, DC8/1 was between 1.3- 3.7-fold more potent than the free antibiotic (Table 2).

**Figure 2.**
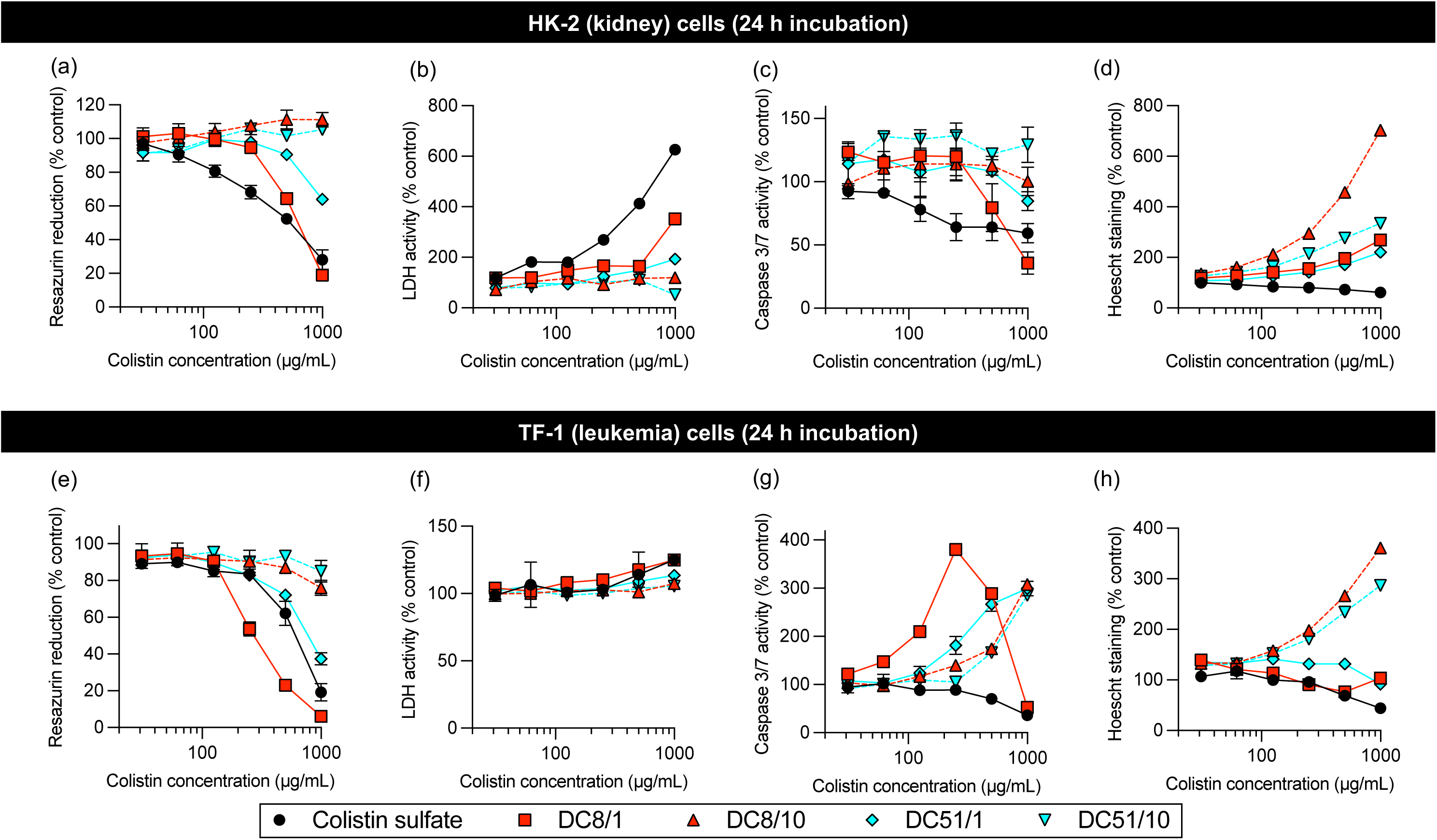
Detection of resazurin reduction (metabolic activity), LDH leakage (cell-membrane integrity, necrosis), caspase 3/7 activity (apoptosis) and Hoechst 33342 staining (DNA content) under multiplex conditions of (a-d) HK-2 and (e-h) TF-1 cells incubated for 24 h with colistin sulfate and dextrin-colistin conjugates. Data represent mean (±1SD, n =3). Where error bars are invisible, they are within size of data points.

Colistin sulfate induced dose-dependent necrosis in all cell lines at 24 and 72 h (Figures 2b,f and S1b,f), with no evidence of caspase 3/7-dependent apoptosis (Figures 2c,g and S1c,g). In contrast, amide-linked conjugates showed dose-dependent caspase 3/7 activity (DC8/1 > DC51/1 > DC8/10 > DC51/10), in all leukemia cell lines at 24 h post-treatment (3.8-fold increase in caspase 3/7 activity in TF-1 cells treated with DC8/1 at 0.25 mg/mL colistin base, Figure 2g). However, by 72 h, no caspase 3/7 activity was seen. No comparable caspase 3/7 upregulation was seen in HK-2 cells for any of the treatments tested (Figures 2c and S1c). Overall, patterns of concentration-dependent cytotoxicity were similar in all leukemia cell lines tested, in contrast to non-cancerous HK-2 cells (Figure 2 and S1).

Differences in concentration-dependent cell viability and apoptosis were observed between amide-and ester-linked dextrin-colistin conjugates (Figure 3). Ester-linked conjugates exhibited a similar effect on cell viability as the amide-linked DC8/1, but the latter caused the greatest increase in caspase 3/7 activity (Figure 3a,b). When cell viability and apoptosis data were plotted against the dextrin content of the respective dextrin-colistin conjugate, the cytotoxic effects became more congruent (Figure S2). DC8/1 remained the most potent amide-linked conjugate, however there was no difference in cytotoxicity between the ester-and amide-linked conjugates. Amide-linked conjugates, however, showed more caspase 3/7 activity at equivalent dextrin concentrations, than the ester-linked conjugates. Annexin V binding occurred within a few hours of treatment and a subsequent peak in necrosis was detected (Figure 3c,d). Ester-linked DC8/1e conjugate caused less apoptosis, but more necrosis, than the amide-linked equivalent (DC8/1). Ester-linked conjugates were not tested in further experiments, due to the high batch-to-batch variability in conjugation efficiency and excessive amounts of unconjugated polymer, compared to amide-linked conjugates.

**Figure 3.**
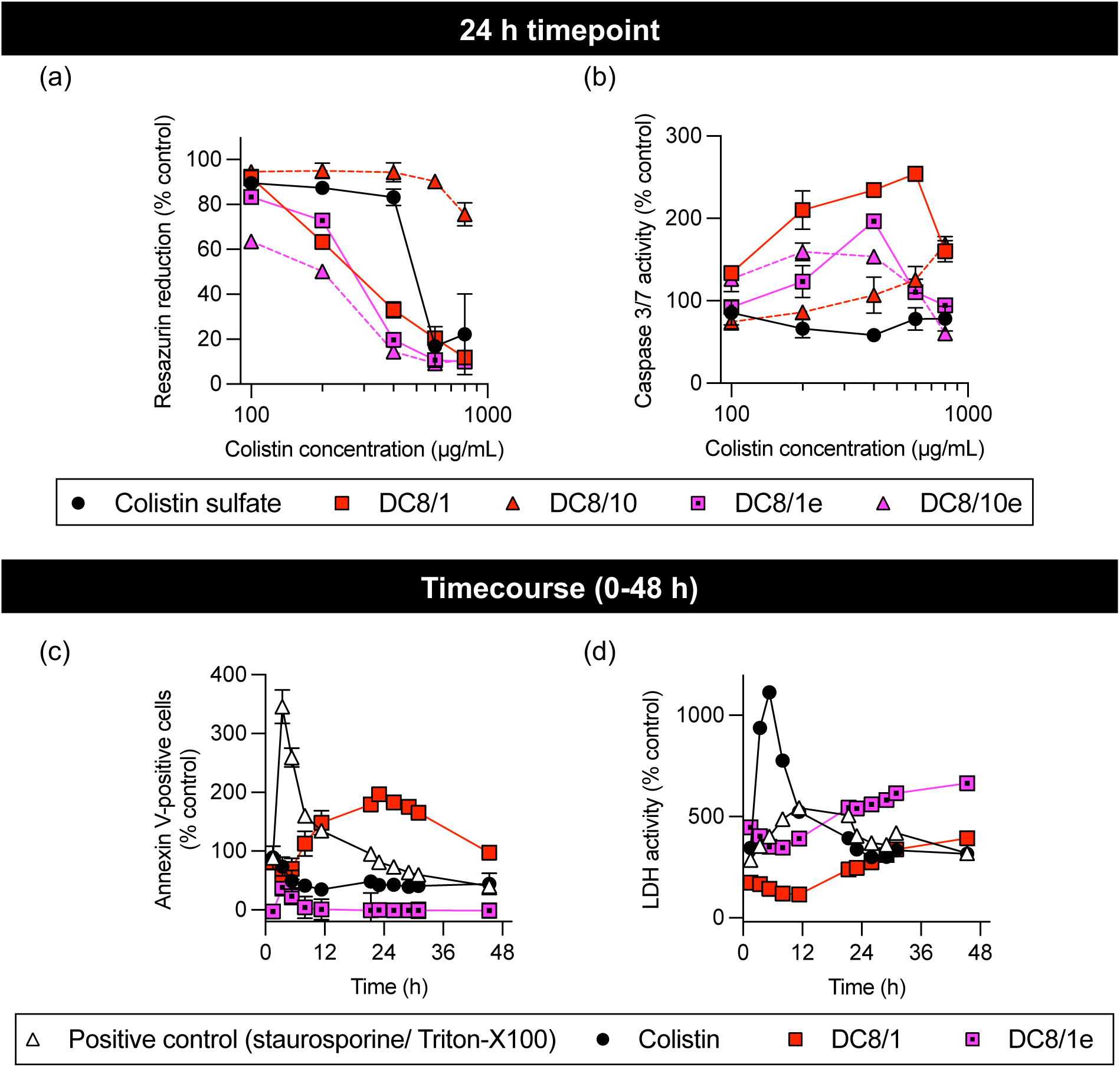
Panels (a) and (b) show detection of resazurin reduction (metabolic activity) and caspase 3/7 activity (apoptosis), respectively, under multiplex conditions of of MV-4-11 cells incubated for 24 h with colistin sulfate and dextrin-colistin conjugates containing amide or ester linkers. Data represent mean (±1SD, n =3). Panels (c) and (d) show time course of annexin V binding and LDH activity (necrosis), respectively, during incubation of MV-4-11 cells with colistin sulfate or ester-and amide-linked dextrin-colistin conjugates at 750 µg/mL colistin base (±1SD, n=3). Where error bars are invisible, they are within size of data points.

Cells treated with DC8/10 and DC51/10 in the multiplex assay exhibited concentration-dependent increases in Hoechst 33342 staining (Figures 2d,h and S1d,h), which suggested DNA accumulation. Subsequent cell cycle analysis using PI failed to show any significant changes in phase distribution associated with different treatments, apart from a limited, but statistically significant, reduction in the G2M population of cells treated with DC8/1, compared to colistin sulfate (Figure 4a).

**Figure 4.**
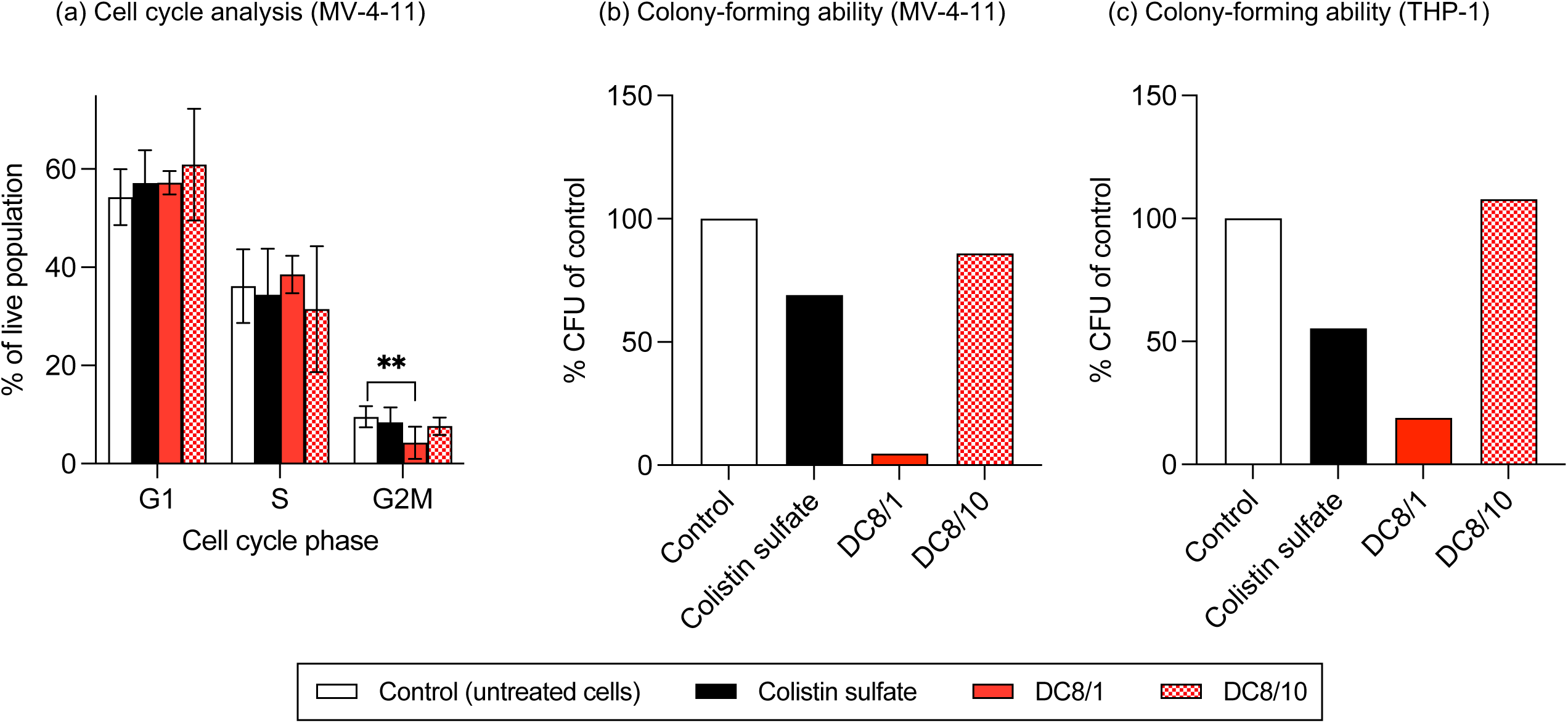
Cell cycle analysis of (a) MV-4-11 cell, and colony-forming ability of (b) MV-4-11 and (c) THP-1 cells following incubation with colistin sulfate or dextrin-colistin conjugates at IC_50_ for 24 h. Data in panel (a) are presented as mean % gated population (±1SD, n=8). Data in panels (b) and (c) are presented as mean % CFU, relative to control (n = 2). ** indicates significance *p*<0.01, compared to untreated control. Where significance is not shown, *p*>0.05 (ns, not significant). Panels (b) and (c) do not show error bars as data is n=2.

### Colony formation

A colony formation assay was employed to determine the potential inhibitory effects of dextrin-colistin conjugates on tumor cell phenotype. Following exposure of leukemia cells to colistin sulfate at IC_50_ for 24 h, colony formation was reduced, compared to control, in both cell lines (MV-4-11: 69±10%, THP-1: 55±1%) (Figure 4b,c). Inhibition of colony formation was most pronounced in cells treated with DC8/1 at the equivalent colistin dose (MV-4-11: 4.7±1.6%, THP-1: 19±2%), while DC8/10 did not alter colony formation in either cell line tested.

### Evaluation of cellular uptake and intracellular fate

To study cellular uptake, AF594-labelled colistin, dextrin and dextrin-colistin conjugates were synthesized, containing <2% free AF594 (Figure S3). The characteristics of all AF594-labelled probes used in these studies are summarized in Table 3.

**Table 3.** Characteristics of AF594-labelled conjugates.

| Compound | Degree of labelling (moles AF594 per mole colistin/ dextrin/ dextrin-colistin conjugates) | Labelling efficiency ($\mu\text{g}$ AF594/mg conjugate) |
| --- | --- | --- |
| Colistin-AF594 | 0.273 | 135.12 |
| Dextrin (8 kDa, 10 mol%)-AF594 | 0.026 | 2.79 |
| DC8/1-AF594 | 0.036 | 2.29 |
| DC8/10-AF594 | 0.092 | 2.11 |

**Abbreviations:** AF594, AlexaFluor 594; $\mu\text{g}$ , microgram; mg, milligram, kDa, kilodalton; DC, dextrin-colistin

Flow cytometry demonstrated that all AF594-labelled conjugates exhibited cell association after 1 h at 37°C (Figures 5 and S4). In all instances, the cell-associated fluorescence was reduced at 4°C, indicating >98% calculated internalization of fluorescent probes. Colistin-AF594 cellular levels were significantly greater than observed for the AF594-labelled dextrin-colistin conjugates. However, DC8/1-AF594 displayed significantly more uptake by MV-4-11 cells than DC8/10-AF594 at 37°C (*p*>0.0001, 2-way ANOVA, Tukey’s multiple comparison test).

**Figure 5.**
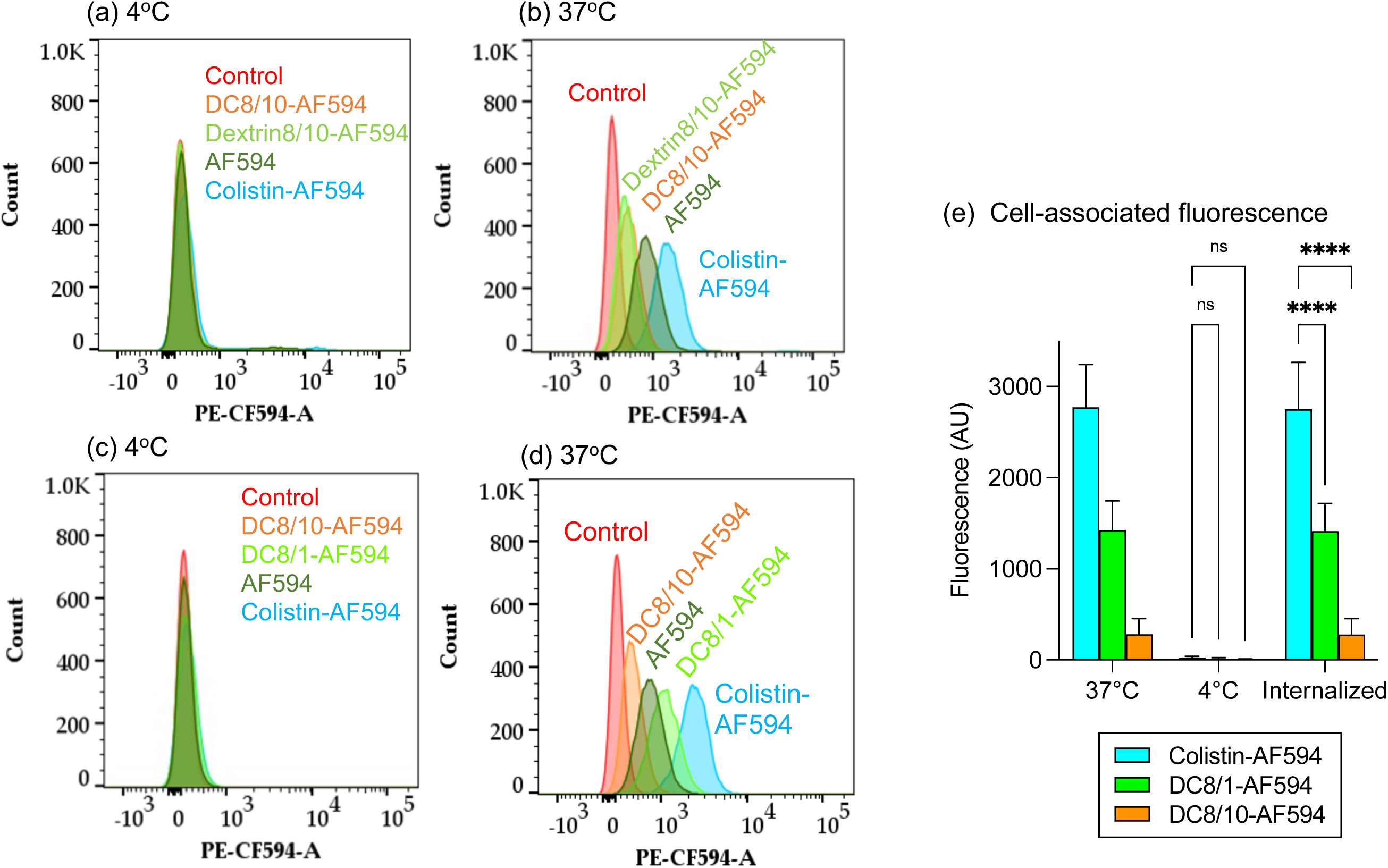
Binding and internalization of AF594 (control) and AF594-labelled colistin, succinoylated dextrin (8k, 10 mol%) and dextrin-colistin conjugates by MV-4-11 cells after 1 h incubation at 4˚C and 37˚C. Panels (a-d) show histograms for cell-associated fluorescence at 4°C ((a) and (c)) and 37°C ((b) and (d)). Panels (a) and (b) show comparative uptake of DC8/10 compared to AF594- labelled colistin and dextrin alone. Panels (c) and (d) show a second experiment comparing the up-take of DC8/1 vs. DC8/10, with appropriate controls. Panel (e) shows cell-associated fluorescence at 37˚C (total association) and 4˚C (external binding) of AF594-labelled colistin, dextrin and dextrin-colistin conjugates (± 1SD, n =). **** indicate significance p<0.0001, compared to colistin-AF594. ± 1SD, n = 6, where * indicates significance *p*<0.05, ** indicates significance *p*<0.01, *** indicates significance *p*<0.001 and **** indicates significance *p*<0.0001 compared to colistin sulfate. Where significance is not shown, *p*>0.05 (ns, not significant).

In line with flow cytometry data, confocal microscopy showed that colistin-AF594 and AF594- labelled dextrin-colistin conjugates were internalized, with punctate labelling resembling endolysosomal structures (Figures 6, 7). There was clear evidence that colistin-AF594 was reaching the lysosomes, manifesting as colocalization with the pulse chased dextran. However, dextrin conjugation significantly reduced colistin internalization, exacerbated by increased succinoylation.

**Figure 6.**
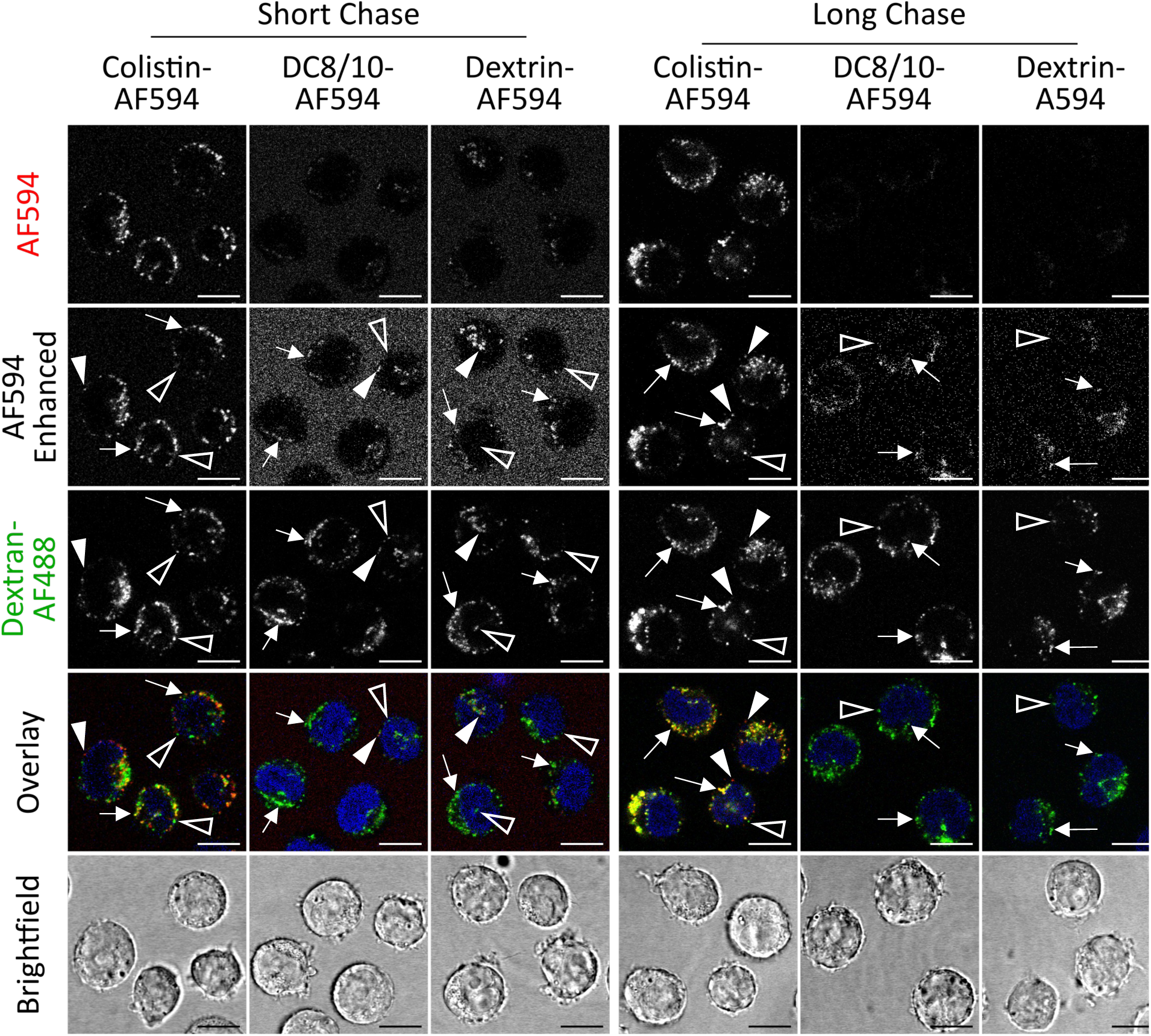
Uptake of Colistin-AF594, DC8/10-AF594 and Dextrin-AF594 (8k/10%) following a 2 h pulse and either 2 h (short) or 16 h (long) chase in MV-4-11 cells. Dextran-AF488 (green) was used to identify late endolysosomes and Hoechst 33342 (blue) was used as a nuclear marker. Scale bar shows 10 µm. Arrows indicate colocalised cargo and lysosomal dextran, solid arrow heads indicate non-colocalised cargo and hollow arrow heads indicate non-colocalised lysosomal dextran. Image intensities have been adjusted so that differences in colocalisation can be more evenly compared. All images are single sections.

**Figure 7.**
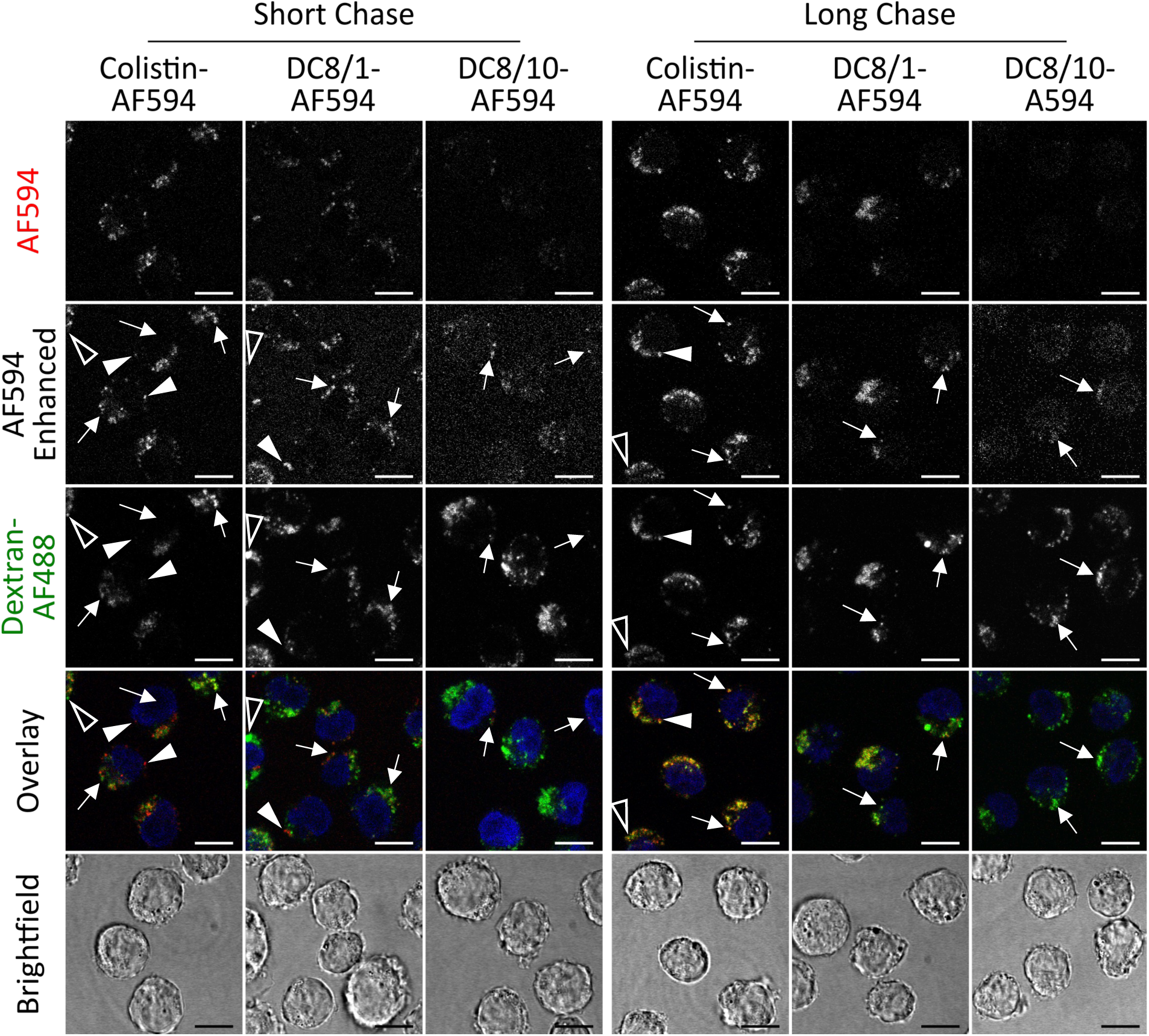
Uptake of colistin-AF594, DC8/1-AF594 and DC8/10-AF594 following a 2 h pulse and either 2 h (short) or 16 h (long) chase in MV-4-11 cells. Dextran-AF488 (green) was used to identify late endolysosomes and Hoechst 33342 (blue) was used as a nuclear marker. Scale bar shows 10 µm. Arrows indicate colocalised cargo and lysosomal dextran, solid arrows indicate non-colocalised cargo and hollow arrow heads indicate non-colocalised lysosomal dextran. Image intensities have been adjusted so that differences in colocalisation can be more evenly compared. All images are single sections.

## Discussion

Targeted therapies, which interfere with specific proteins involved in tumorigenesis, survival and growth, are promising precision medicines for cancer.^35^ The clinical development of targeted therapies has yielded few FDA-approved treatments for AML,^36^ in part, due to unacceptable side effects and an inability to reach their intracellular targets. Drug delivery strategies, such as polymer therapeutics, offer the ability to overcome these issues.^37^ LSD1/KDM1A is overexpressed in many cancers, including AML and has a significant role in regulating differentiation, proliferation and invasion of cancer cells, making it an attractive target for AML treatment.^38^ Speranzini *et al*^14^ reported non-covalent inhibition of LSD1/KDM1A by polymyxin antibiotics, which supported a serendipitous observation by Vertsovsek *et al*,^13^ that polymyxin B was cytotoxic to a range of cancer cell lines, including the human acute lymphocytic leukemia cell line, REH (IC_50_ = _≥_20 µg/mL).

Here, we showed that colistin sulfate was less cytotoxic to leukemia cell lines than a normal kidney cell line. When Speranzini *et al*^14^ treated MV-4-11 cells with colistin, they did not observe any significant alteration in cell growth or H3-Lys^4^/H3-Lys^9^ methylation. However, this study used a relatively low concentration of colistin sulfate (1.2 µg/mL, 0.85 µM), given that our study found an IC_50_ of this agent in the same cells to be 888 µg/mL (768.5 µM). In contrast, other LSD1 inhibitors have shown substantially higher activity towards AML cell lines, for example, tranylcypromine-based compounds developed by Fioravanti *et al* had IC_50_ values of 0.4 and 2.5 µM in MV-4-11 cells (48 h incubation)^39^ and the LSD1 inhibitor, IMG-7289 had an IC_50_ values of 0.007 µM in the same cell line (48 h incubation).^40^

Speranzini *et al* hypothesized that colistin was unable to cross the plasma membrane efficiently, however, we showed here that fluorescently-labelled colistin was rapidly endocytosed by all 3 leukemia cell lines tested, with compelling evidence of further traffic to lysosomes. Contrary to our initial hypothesis, however, dextrin conjugation reduced internalization of colistin by leukemia cells, but did not alter its intracellular localization or promote nuclear localization.

Although all leukemia cell lines tested had similar mRNA expression of KDM1A mRNA expression (summarized in Table S1)^41^, the relative cytotoxicity of colistin sulfate or dextrin-colistin conjugates varied between the leukemia cell lines tested; MV-4-11 showing the largest reduction in IC_50_ for DC8/1 (4-fold lower). Cellular uptake of fluorescently-labelled compounds were similar for all three leukemia cell lines tested, suggesting that inconsistent internalization across the cell lines was not the cause of sensitivity differences.

When dextrin was conjugated to colistin, variable effects on cytotoxicity were observed, which were dependent on dextrin chain length, degree of succinoylation and type of linker used. The most potent effect seen in leukemia cell lines was from dextrin-colistin conjugates containing 8,000 g/mol dextrin, with 1 mol% succinoylation and attached to colistin by an amide bond (DC8/1). Although ester-linked conjugates showed similar potency, when their effects were compared to dextrin content of the conjugate, they caused less apoptosis than the amide-linked conjugates. Varache *et al*^42^ have previously demonstrated that increasing the degree of dextrin succinoylation typically leads to a greater number of linkers bound to colistin; corresponding to the conjugate’s antimicrobial activity. Even after degradation by amylase, residual linker groups, with differing lengths of glucose units attached, prevent complete restoration of colistin activity.

Compared to the amide-linked dextrin-colistin conjugates, colistin sulfate and ester-linked dextrin-colistin conjugates induced less annexin V binding and caspase 3/7 dependent cell death. This suggests that the residual linker groups remaining attached to colistin may play a key role in the activation of caspase 3/7 apoptosis.^27^ Conjugation efficiency of ester-linked conjugates was generally lower than those containing amide linkers, meaning that conjugates contained more unreacted dextrin. The presence of free dextrin in polymer conjugates has been previously reported.^43^ Removal of unreacted dextrin using size exclusion chromatography is challenging, since its size is relatively similar to the dextrin-colistin conjugate. At the concentrations used for antibacterial activity (typically 2-16 µg colistin base/mL), the succinoylated dextrin concentration contained in the conjugates is not toxic. However, to achieve the high concentrations of colistin required to cause cell death in leukemia cells lines, the dextrin content of conjugates exceeded safe levels and masked the effects of colistin.

The mechanism of polymyxin nephrotoxicity has been widely studied and is related to colistin’s interaction with oxidative stress and death-related pathways such as necrosis, apoptosis and autophagy.^44^ Here, necrosis, but not apoptosis, was the main mode of cell death by colistin sulfate in both, kidney and leukemia cell lines. However, DC8/1 caused caspase-dependent apoptosis in leukemia cells, suggesting that this chemical modification of colistin alters the mechanism of cell death. Induction of annexin V binding, and thus apoptosis, occurred within a few hours of exposure to DC8/1, and cell death was extensive by 24 h, suggesting that the mechanism of cytotoxicity may not be mediated through induction of differentiation, for example, by LSD1 inhibition. Preliminary studies found no evidence of upregulation of CD11b or CD86 following 48 h treatment with DC8/1 and 24 h recovery (data not shown).

Whilst at higher concentrations, some of the dextrin-colistin conjugates caused an apparent increase in DNA content, subsequent analysis did not show any cell cycle disruption. As a characterized efflux pump substrate (e.g. ATP-binding cassette (ABC) transporter proteins) we propose that the increased Hoechst 33342 uptake that was greatest for dextrin-colistin conjugates containing 10 mol% succinoylated dextrin, may be due to altered activity or expression of these proteins in the leukemia cell lines, rather than G2 cell cycle accumulation. Cell cycle disruption has been linked to LSD1/KDM1A inhibition, for example, Nicosia *et al*^45^ demonstrated cell cycle inhibition, via downregulation of a LSD1/KDM1A target in the acute promyelocytic leukemia cell line, NB-4. Interestingly, although none of the colistin treatments tested here altered the cell cycle in leukemia cells, Dai *et al*^44^ did observe cell cycle arrest in a murine nephrotoxicity model and Eadon *et al* found that homogenized kidney tissue of mice who were administered colistin had altered expression of genes that regulate the cell cycle,^46^ which suggests a tissue-specific effect. Reduced colony forming ability of leukemia cell lines correlated with cell viability at 24 h, suggesting that colony forming cells were not preferentially targeted by either colistin sulfate or the dextrin-colistin conjugates.

Here, we were unable to establish the exact mechanism of enhanced cytotoxicity of DC8/1 and we did not observe any evidence of LSD1/KDM1A inhibition. However, Smitheman^47^ observed peak LSD1/KDM1A inhibition of leukemia cell lines (including MV-4-11 and THP-1) treated with sub-lethal concentrations of LSD1/KDM1A inhibitors (GSK2879552 and GSK-LSD1) at 3-6 days, with no evidence of apoptosis. Colony formation results suggested that the main impact on functionality of surviving cells occurred within 24 h of treatment, which supports the timescales used in our cytotoxicity experiments.

### Clinical potential of dextrin-colistin conjugates to treat AML

Drug repurposing (also known as drug repositioning, reprofiling or re-tasking) is a process of identifying new uses for existing drugs beyond the scope of the original medical indication.^48^ The advantages of this approach include lower risk of failure, shorter time to market and reduced drug development costs.^49^ Recently, several antibiotics have been repurposed to treat cancer.^50^ For example, levofloxacin was shown to inhibit proliferation and induce apoptosis of lung cancer cells in preclinical studies^51^ and doxycycline has shown anti-proliferative and pro-apoptotic activity in cervical cancer and breast cancer cell lines.^52,53^ Of the 371 drugs currently being evaluated in clinical trials as potential treatments for various cancers, 10 are antibiotics.^54^ Nevertheless, antibiotic repurposing remains controversial, due to the potential impact on the development of antimicrobial resistance (AMR),^55^ which correlates strongly with increased consumption.^56,57^ AML is an aggressive condition that requires rapid and effective treatment that can last for several years. Prolonged antibiotic use may induce drug resistance among gut microbes or create antibiotic resistance that could reduce the effectiveness of treatments for common infections or render the patients more prone to infection.

When intravenous colistin (usually administered as the prodrug, colistimethate sodium (CMS)) is used to treat patients with Gram-negative bacterial infections, a target average colistin plasma concentration at steady-state (C_ss,avg_) of 2 mg/L is usually used,^58^ corresponding to the minimum inhibitory concentration clinical breakpoint for several Gram-negative pathogens. However, at plasma colistin concentrations of >2.5 mg/L, the risk of concentration-dependent nephrotoxicity increases substantially.^59,60^ In agreement with previous studies,^11^ dextrin-colistin conjugates were ∼3-fold less toxic to the kidney cell line. While this is a key development in tackling the cytotoxicity of colistin to normal tissues, ultimately, it is probably unfeasible that the IC_50_ values seen for colistin and dextrin-colistin conjugates in leukemia cell lines (exceeding 235 mg/L) could be achieved clinically without causing severe nephrotoxicity. Although therapeutic concentrations of dextrin-colistin conjugate are unlikely to be achieved clinically, this approach could be applied to modify other, more potent, non-antibiotic therapeutics that have shown promise for the treatment of AML to reduce toxic side effects and boost potency.

## Conclusion

This study has demonstrated a potentiation of colistin’s anti-leukemic activity by covalent, irreversible attachment of dextrin, which was unlikely to be dependent on LSD1/KDM1A. Our results show that, despite cellular uptake being effectively reduced by dextrin attachment, polymer conjugation can enhance the biological activity of a drug, alter its mechanism of action and localize drugs to endolysosomes. Whilst clinical translation of dextrin-colistin conjugates for the treatment of AML is unlikely due to the potential to promote AMR and the relatively high colistin concentrations required for anticancer activity *in vivo*, the ability to potentiate the effectiveness of an anticancer drug by polymer conjugation, while reducing side effects and improving biodistribution of the drug, is very attractive, and this approach warrants further investigation.

## Acknowledgments

This article is dedicated to the memory of Dr Konrad Beck, who presented the idea of investigating dextrin-colistin conjugates as an LSD1 inhibitor to treat cancer. The authors acknowledge Miss Gilda Pourbahram and Miss Amanda Gilbert for their contributions to literature review and method development, respectively. We also acknowledge the technical cell culture support provided by Dr Sarah Youde and Dr Maria Stack and thank Dr Catherine Naseriyan from Central Biotechnology Services, Cardiff University for their technical expertise in FACS.

## Supplementary materials

The following supporting information can be downloaded at:

## Data Sharing Statement

All the data contained in the article is available upon reasonable request.

## Author Contributions

Conceptualization, S.R., A.T.J., A.T., D.W.T. and E.L.F.; methodology, S.R., M.V., E.J.S, A.T.J., A.T., D.W.T. and E.L.F.; formal analysis, S.R., M.V., E.J.S, A.T.J., A.T., D.W.T. and E.L.F.; investigation, S.R, M.V., E.J.S., G.P., E.L.F.; resources, A.T.J., A.T., D.W.T., and E.L.F.; data curation, S.R, M.V., E.J.S., E.L.F.; writing—original draft preparation, S.R. and E.L.F.; writing— review and editing, S.R., M.V., E.J.S, A.T.J., A.T., D.W.T. and E.L.F.; visualization, S.R., M.V., E.J.S. and E.L.F.; supervision, A.T.J., A.T., D.W.T. and E.L.F; project administration, S.R. and E.L.F.; funding acquisition, S.R., A.T.J., A.T., and E.L.F. All authors have read and agreed to the published version of the manuscript.

