## Supplementary figures and tables for "Modification of the antibiotic, colistin, with dextrin causes enhanced cytotoxicity and triggers apoptosis in myeloid leukemia"

Rizzo *et al*

**Table S1.** Characteristics of cell lines used in this study.

| Cell line | Source | Patient | Pathology | Culture medium | KDM1A mRNA expression (RNAseq)* |
|---|---|---|---|---|---|
| MV-4-11 | Peripheral blood | Caucasian male, 10yrs | biphenotypic B myelomonocytic leukemia | IMDM withGlutaMAX™, 10% v/v heat-inactivated (HI)-FBS | 5.08 |
| THP-1 | Peripheral blood | Japanese male, 1yr | acute monocytic leukemia | RPMI 1640 withGlutaMAX™, 10% HI-FBS | 5.40 |
| TF-1 | Bone marrow | Japanese male, 35yrs | erythroleukemia | RPMI-1640, 10% v/v HI-FBS, 2ng/mL rhGM-CSF | 6.11 |
| HK-2 | Kidney proximal tubule | Caucasian male, adult | Normal | K-SFM with L-glutamine, epidermal growth factor, bovine pituitary extract | N.D. |

*from Barretina *et al.*[38] RNAseq TPM gene expression data for just protein coding genes using RSEM. Log2 transformed, using a pseudo-count of 1.
N.D., not determined

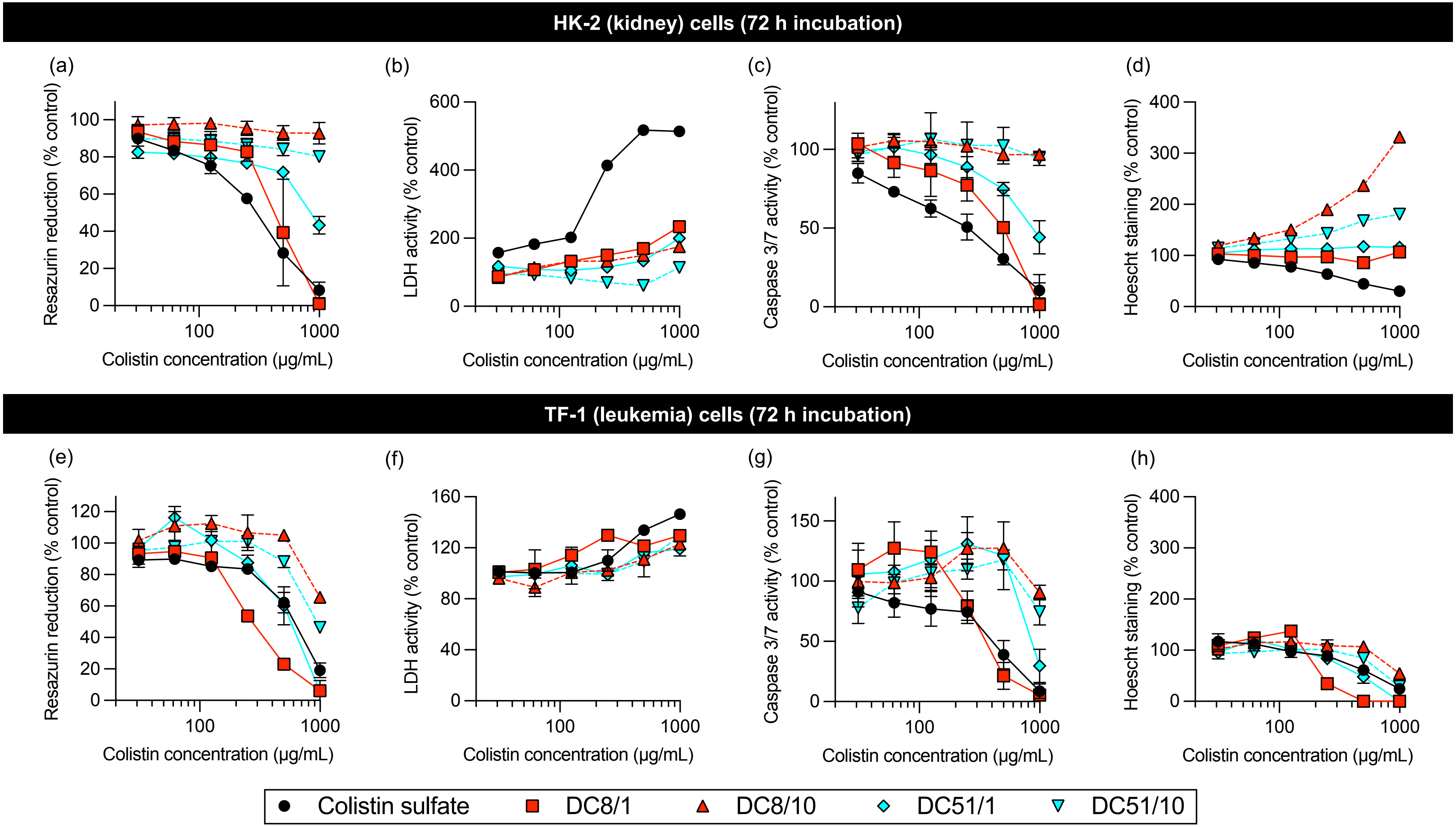


**Figure S1.** Detection of resazurin reduction (metabolic activity), LDH leakage (cell-membrane integrity, necrosis), caspase 3/7 activity (apoptosis) and Hoechst 33342 staining (DNA content) under multiplex conditions of (a-d) HK-2 and (e-h) TF-1 cells incubated for 72 h with colistin sulfate and dextrin-colistin conjugates. Data represent mean (±1SD, n =3). Where error bars are invisible, they are within size of data points.

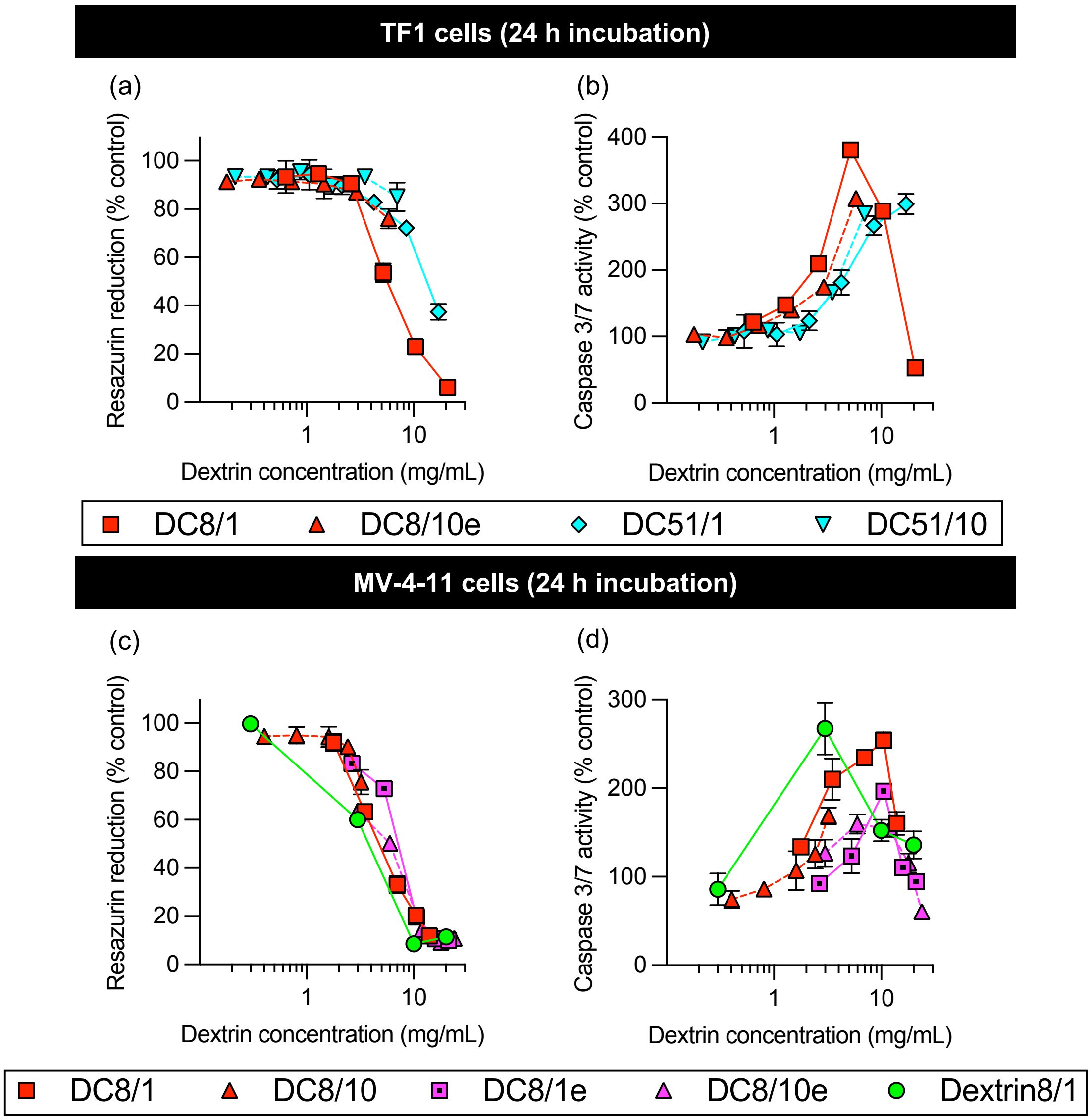


**Figure S2.** Detection of resazurin reduction (metabolic activity) (panels (a) and (c)) and caspase 3/7 activity (apoptosis) (panels (b) and (d)) under multiplex conditions of MV-4-11 cells incubated for 24 h with colistin sulfate and dextrin-colistin conjugates containing amide or ester linkers. Data represent mean (±1SD, n =3). Where error bars are invisible, they are within size of data points.

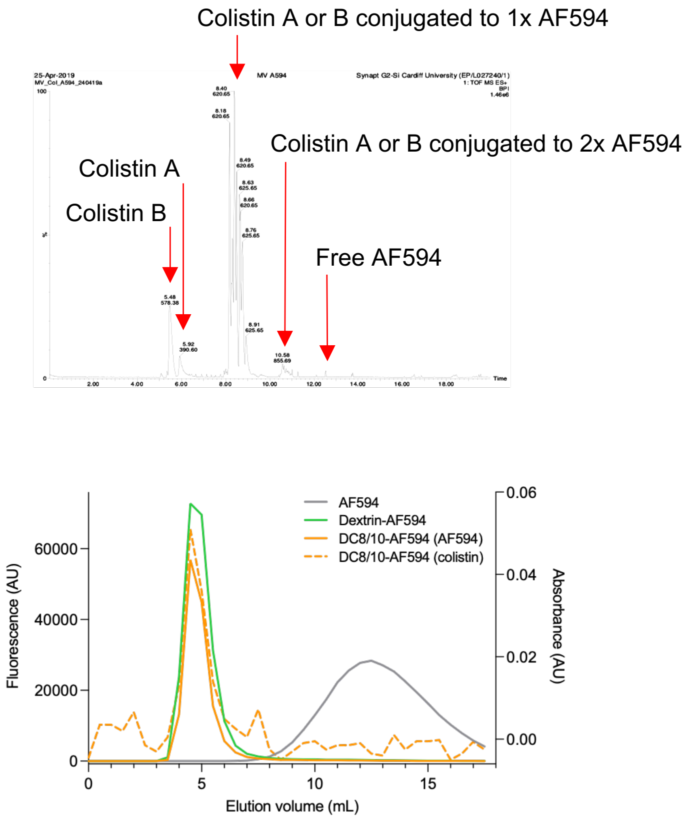


**Figure S3.** Representative characterization of AF594-labelled conjugates using (a) LC-MS and (b) PD-10 column separation and analysis of fractions using fluorescence (AF594 content) and absorbance (colistin content).

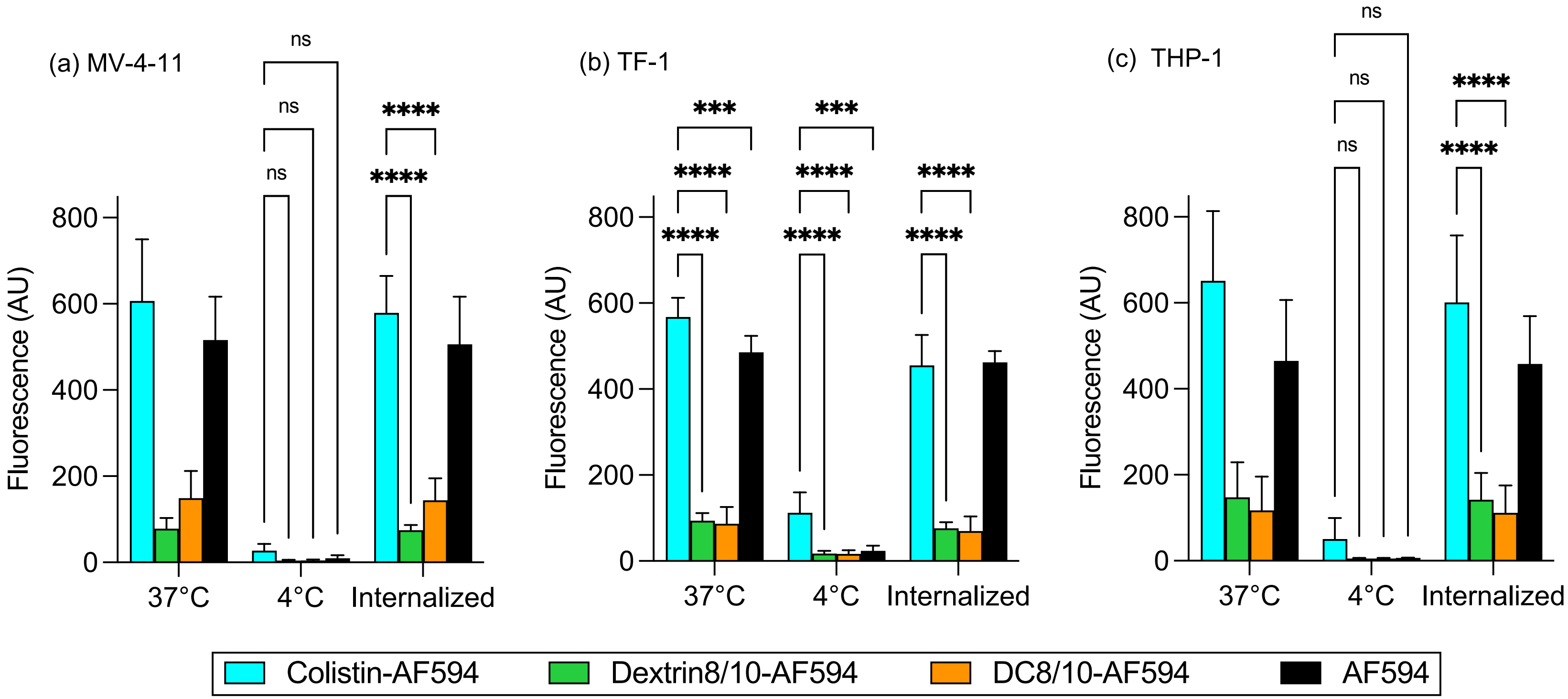


**Figure S4.** Cell-associated fluorescence at 37˚C (total association) and 4˚C (external binding) of AF594-labelled colistin, dextrin and dextrin-colistin conjugate (DC8/10) by (a) MV-4-11, (b) TF-1 and (c) THP-1 cells after 1 h incubation at 4 and 37˚C (±1SD, n = 5 to 8), where * indicates significance $p$<0.05, ** indicates significance $p$<0.01, *** indicates significance $p$<0.001 and **** indicates significance $p$<0.0001 compared to colistin-AF594. Where significance is not shown, $p$>0.05 (*ns*, not significant).
